# Elevated levels of interleukin-27 in early life compromise protective immunity during neonatal sepsis

**DOI:** 10.1101/777839

**Authors:** Brittany G. Seman, Jordan K. Vance, Travis W. Rawson, Michelle R. Witt, Annalisa B. Huckaby, Jessica M. Povroznik, Shelby D. Bradford, Mariette Barbier, Cory M. Robinson

## Abstract

Neonates are at increased risk for bacterial sepsis as a result of immature immunity. We established that the immune suppressive cytokine interleukin (IL)-27 is elevated in early life. In the present work, we hypothesized that increased levels of IL-27 may predispose the neonatal population to more severe infection during sepsis. In a neonatal sepsis model, systemic IL-27 levels continued to rise during infection. Peripheral tissue analysis revealed systemic IL-27 expression, while myeloid cell profiling identified Gr-1 and F4/80-expressing cells as the most abundant producers of IL-27 during infection. Increased IL-27 levels were consistent with increased mortality that was improved in WSX-1^-/-^ mice that lack a functional IL-27 receptor. Infected WSX-1^-/-^ pups exhibited improved weight gain and reduced morbidity. IL-27 signaling in WT mice promoted increased bacterial burdens and systemic inflammation compared to WSX-1^-/-^ neonates. This was consistent with more efficient bacterial killing by Ly6B.2^+^ myeloid cells and macrophages from WSX-1-deficient compared to wild-type neonates. Live animal imaging further supported a more severe and disseminated infection in WT neonates. This is the first report to describe the impact of elevated early life IL-27 on the host response in neonates while also defining the cell and tissue sources of cytokine. IL-27 is frequently associated with suppressed inflammation. In contrast, our findings demonstrate that IL-27 promotes inflammation during neonatal sepsis by directly compromising control of bacteria that drive the inflammatory response. Collectively, our results suggest that IL-27 represents a therapeutic target to limit susceptibility and improve infectious outcomes in neonatal sepsis.

**IMPORTANCE:** A number of differences in the neonatal immune response compared with adults have been well described. However, a mechanistic understanding of what needs to be overcome in the neonate to generate a more protective immune response during acute bacterial infection has been limited. The work described here helps fill the gap of what is necessary to overcome in order to achieve improved host response to infection. To further the novelty, IL-27 has not previously been attributed to dysfunction or deficiency in neonatal immunity. Our results enhance the understanding of IL-27 biology in the neonatal population while providing evidence that elevated IL-27 levels limit a protective immune response and are detrimental during neonatal sepsis. Strategies aimed at targeting circulating IL-27 concentrations early in life have the potential to improve control of bacterial infection in neonates.

## INTRODUCTION

Neonates are highly vulnerable to bacterial infections and at increased risk of mortality. Accuracy identifying the true global incidence of neonatal sepsis is influenced by challenges with diagnostic criteria and reliable reporting, but current estimates indicate approximately 3 million infections annually (1). In the United States alone, greater than 75,000 neonatal infections are reported annually due to sepsis (2). The rate of cases per live birth increases considerably with factors such as low-birth-weight and premature delivery (2). Sepsis is a leading cause of morbidity mortality among infants in the first few days of life at any birth weight (3). This is especially true for very low birth weight infants where sepsis also significantly increases the length of hospital stay (4, 5).

Increased susceptibility to infection in neonates is reflective of a distinct immune profile relative to adults that is often referred to as immature. Phenotypic and functional differences in innate and adaptive immune function have been described in early life. In general, neonatal immunity is considered biased toward a Th2/Treg response (6). Fewer numbers of immune cells have been found in circulation with defects reported in microbial elimination processes, antigen presentation, T cell priming, and T cell receptor repertoires (7-9). In addition, increased amounts of cytokines such as IL-10, IL-23, and IL-27 are present, further supporting an anti-inflammatory bias (10-12). This is consistent with reduced production of tumor necrosis factor (TNF)-α from neonatal cells in response to TLR ligands compared with those from adults (13). Since adequate Th1 responses can be induced in neonates *in vivo* when given the appropriate stimulus, innate immune cells may provide signals that delay or misdirect the adaptive immune response (14). Cumulatively, these immune findings are thought to contribute to the increased susceptibility of neonates to infection.

Interleukin-27 (IL-27) is a heterodimeric cytokine that consists of the Epstein-Barr virus-induced gene 3 (EBI3) and IL-27p28 proteins (15). Engagement of the IL-27 receptor, composed of WSX-1 and gp130, predominantly activates JAK-STAT signaling (16-18). IL-27, similar to other members of the IL-12 family, was originally described as a cytokine that could drive proliferation of naïve CD4^+^ T cells (19). However, mice deficient for WSX-1 mount Th1 responses (18, 20-23). In these same animals, models of chronic disease and infection demonstrate T cell hyperactivity suggesting that additional immune suppressive activity may dominate (18, 21-24). Indeed, IL-27 antagonizes inflammatory T cell subsets by blocking IL-2 production, and activates IL-10 production by Treg cells (25). Similarly, innate immune function is inhibited by IL-27. In macrophages, inflammatory cytokine production, inflammatory cytokine receptor expression and signaling, intracellular trafficking to lysosomes, and lysosomal acidification are negatively regulated by IL-27 (22, 26-31). This regulatory activity has been shown to compromise control of *M. tuberculosis, S. aureus, P. aeruginosa*, and *E. coli* (12, 26, 27, 30, 31). On the other side of the spectrum, IL-27 induces an inflammatory profile in monocytes (32). Cumulatively, this body of literature suggests that IL-27 has important immune regulatory function and opposes clearance of bacteria by macrophages. The effect of IL-27 may be cell type and context-dependent, and the net influence on the compete host response in neonates has not been understood.

We have established that IL-27 is produced at elevated levels early in life. Human macrophages derived from umbilical cord blood express IL-27p28 and EBI3 genes at increased levels compared with macrophages derived from adult peripheral blood (11). This was accompanied by greater levels of secreted IL-27 protein (33). Similarly, transcript levels for IL-27 genes were increased in the spleens of neonatal and infant mice relative to adults (11). A similar pattern of IL-27 production was observed in splenic macrophages from neonatal and infant mice (11). Recently, myeloid-derived suppressor cells (MDSCs) were shown to be a significant source of IL-27, and these cells were more abundant in neonates than other age groups (12). Other reports have shown a greater abundance of MDSCs in human blood during the neonatal period (34, 35).

Considering the immune suppressive activity of IL-27, we have hypothesized that elevated IL-27 early in life contributes to enhanced susceptibility to bacterial infection. IL-27 has been suggested as a biomarker for critically ill children and more recently declared a biomarker for early onset neonatal sepsis (EONS) (36, 37). In the present body of work, we examined the impact of IL-27 on host protection during neonatal sepsis. We developed a murine model of EONS in response to *Escherichia coli*. While Group B streptococci are the leading cause of EONS overall, *E. coli* is responsible for the majority of deaths and is the leading cause when pre-term and very-low birth weight babies are considered independently (3, 38). Our findings demonstrate that IL-27 compromises host control of bacteria, consistent with elevated levels of inflammation and increased mortality.

## MATERIALS AND METHODS

### Animals

Breeding pairs of C57BL/6 (WT) or WSX-1-deficient (KO) mice on a C57BL/6 genetic background were purchased from Jackson Laboratories (Bar Harbor, ME) and maintained at West Virginia University School of Medicine. Male and female pups were used for experimental infection. Mice in this study were defined as neonates through 8 days of life as described previously (11, 12). Blood and tissues were collected from mice at the appropriate age by humane procedures. All procedures were approved by the West Virginia University Institutional Animal Care and Use Committees and conducted in accordance with the recommendations from the *Guide for the Care and Use of Laboratory Animals* by the National Research Council (NRC, 2011).

### Bioluminescent E. coli

*E. coli* O1:K1:H7 (ATCC, Manassas, VA) was transformed with the pGEN-luxCDABE plasmid (Addgene #44918) by electroporation using a Micropulser (Bio-Rad, Hercules, CA). This plasmid contains five *lux* genes with a selectable ampicillin resistance marker (Amp^R^). To generate a stable integration, *E. coli* O1:K1:H7 was transformed with the pMQ-tn-PnptII-lux suicide vector (a gift by Dr. Robert Shanks, University of Pittsburgh). This plasmid contains a transposable element upstream of five *lux* genes, with a selectable ampicillin resistance marker (AmpR). Transformation was performed by mating with the auxotrophic strain RHO3 (39). Transformants were selected on ampicillin-supplemented LB agar and screened for luminescent signal on a chemiluminescent imager. Luminescence was monitored through 48 h of growth and infection to assess plasmid retention. Intravital imaging was performed with *E. coli* expressing luciferase from the transformed plasmid. *E. coli* with stably integrated *lux* genes were used in gentamicin protection assays to evaluate bacterial clearance.

### Murine sepsis infection model

Neonatal pups at the ages of 3 or 4 days were infected subcutaneously in the scapular region with *E. coli* strain O1:K1:H7. The bacteria from pre-titered frozen cultures were washed with PBS, centrifuged at 2,000 x g for 5 min, and suspended in a volume of PBS equivalent to an inoculum of 50 μL/mouse. Mice were inoculated using a 28-gauge insulin needle (Covidien, Dublin, Ireland). Vehicle (PBS)-inoculated pups were identified from bacteria-infected pups using a tail snip. Survival studies were performed with an inoculum of ∼10^7^ CFUs/mouse representing an approximate LD_50_. Other experiments to evaluate infection-related parameters were performed with a target inoculum of ∼2 × 10^6^ CFUs/mouse to reduce mortality so that sufficient numbers of control and infected animals could be studied in each experiment. Weights of mice were recorded immediately prior to infection and then daily thereafter. Following infection, mice were monitored twice daily for signs of morbidity. Mice exhibiting signs of morbidity (i.e., unable to right themselves, significant weight loss, and lack of movement) that met endpoint criteria were humanely euthanized. Peripheral tissues (spleen, liver, kidney, brain, and lung) isolated from pups were placed in PBS on ice. Blood was deposited in tubes that contained 5 μL of 500 mM ethylenediamine tetraacetate acid (EDTA, Fisher Scientific, Fair Lawn, NJ).

### Bacterial burdens

Peripheral tissues were homogenized in PBS using a handheld pestle motor (Kimble Chase, Vineland, NJ). Tissue homogenates and blood were serially diluted in PBS and bacteria enumerated by standard plate counting on tryptic soy agar (TSA; Becton, Dickinson and Company, Sparks, MD). Agar plates were incubated at 37°C overnight.

### Intracellular Cytokine Staining

Spleens were crushed in a 40-µm nylon strainer. Single cell suspensions were treated with ACK lysis buffer (Lonza, Walkersville, MD) to lyse red blood cells and washed in PBS that contained 10% FBS. Blood was pooled from control or infected mice and washed in PBS. Peripheral blood mononuclear cells (PBMCs) were isolated by Ficoll (GE Healthcare Life Sciences, Chicago, IL) density gradient centrifugation at 400 x g for 30 minutes. Splenocytes and PBMCs were then treated with FcR blocking reagent (Miltenyi Biotec, Bergisch Gladbach, Germany) and GolgiStop (Protein Transport Inhibitor, Becton Dickinson, Franklin, NJ) to inhibit protein secretion. Cell surface markers were immunolabeled with anti-Gr-1 (PE Rat Anti-Mouse Ly6G and Ly6C, BD Pharmingen, Franklin, NJ), F4/80 (Anti-F4/80 PE, Miltenyi Biotec), CD11c (CD11c-PE, Miltenyi Biotec), or CD115 (CD115-PE, Miltenyi Biotec), washed, and fixed with 3% paraformaldehyde. Intracellular cytokine IL-27 was labeled with anti-mIL-27 (R&D Systems, Minneapolis, MN) as described previously (11). Immunolabeled cells were analyzed with a BDFortessa flow cytometer and FCS Express (version 6; De Novo Software, Glendale, CA).

### Quantitative real time PCR

Spleens were homogenized in TRI Reagent^®^ (Molecular Research Center, Cincinnati, OH). Using the commercial product protocol, the upper aqueous layer following phase separation was mixed with an equal volume of 75% ethanol and transferred to EZNA RNA isolation columns (Omega Biotek, Norcross, GA). iScript(tm) cDNA synthesis reagents (Bio-Rad, Hercules, CA) were used to generate first strand cDNA according to manufacturer protocol. Real time cycling of reactions that included cDNA diluted 15-fold from above, gene-specific primer probe sets (Applied Biosystems, Foster City, CA), and iQ^™^ Supermix (Bio-Rad) was performed in triplicate using a Step One Plus (Applied Biosystems) real time detection system. Gene-specific amplification was normalized to β-actin as an internal reference gene. Log_2_ transformed changes in gene expression relative to control spleens were determined using the formula 2^-ΔΔCt^.

### Cytokine measurements

Blood was collected from mice during euthanasia at the indicated age and serum collected by standard techniques. IL-1, IL-6, and TNF-α serum levels were measured using multiplexed luminescent detection reagents according to manufacturer’s protocol (MesoScale Discovery, MSD, Rockville, MD). Serum or culture supernatant concentrations of IL-27p28 were measured using single analyte luminescent detection reagents (MesoScale Discovery). Results were analyzed using MSD Discovery Workbench software. Protein standards were assayed in parallel with samples.

### In vivo imaging

Neonatal pups were imaged using an IVIS SpectrumCT (Perkin Elmer, Waltham, MA). Mice were infected with the bioluminescent *E. coli* and imaged over time for location and intensity of luminescence. To decipher between individual mice over time, tails were tattooed using a 28-gauge insulin needle that inserts green or black tattoo paste (Ketchum Manufacturing, Lake Luzerne, NY). Pups were anesthetized using an isoflurane chamber and kept under anesthesia during imaging. Luminescence signal and images were processed using Living Image 4.5 software (Perkin Elmer, Waltham, MA). Briefly, signal was quantified by region of interest (ROI) construction around the area of luminescence in 2D images. Signal was quantified in radiance units, and represented as total flux (photons/second). Luminescence scales were set according to the colorimetric scale for WT or KO mice at each time point due to the profound differences in signal between the two genotypes.

### Gentamicin protection bacterial clearance assays

To generate bone marrow-derived macrophages (BMDM), bone marrow progenitors were extracted from femurs, tibias, radius-ulna, and humerus bones of C57BL/6 or WSX-1^-/-^ neonatal mice in α-MEM (MEM; Corning, NY) containing 10% FBS, 2 mM glutamine, and 100 U/mL penicillin/streptomycin. The contents were strained through a 40-um nylon strainer to remove residual tendon and ligament tissue, and erythrocytes were lysed using 0.2% sodium chloride and neutralized in 1.6% sodium chloride. Following a wash with phosphate-buffered saline (PBS), bone marrow cells were differentiated in DMEM that contained 2 mM glutamine, 25 mM HEPES, 10% FCS, and 10% L-cell-conditioned medium for 5-7 days at 37°C with 5% CO_2_ as described previously (12). MDSCs were cultured in DMEM that contained 2 mM glutamine, 25 mM HEPES, and 10% FCS. Ly6B.2^+^ and F4/80^+^ cells were isolated from splenocytes prepared as described above by immunomagnetic selection using Miltenyi isolation reagents (Miltenyi Biotec, Bergisch Gladbach). BMDMs, F4/80^+^, or Ly6B.2^+^ cells, were cultured with luciferase-expressing *E. coli* at a multiplicity of infection (MOI) of 50 for 1 h at 37°C and 5% CO_2_. The medium was then replaced with fresh supplemented with gentamicin (100 μg/mL) and the cultures returned to incubation for an additional 4 h. Bacterial luminescence was measured using a Molecular Devices iC3 at 3 and 6 h post-infection (San Jose, CA).

### Statistical Analysis

All statistical analyses were performed using GraphPad Prism software (version 8; La Jolla, CA). Data was tested using the appropriate parametric or nonparametric measures, as indicated in the figure legends. The threshold for statistical significance was set to 0.05.

## RESULTS

### IL-27 levels rise during neonatal sepsis

Neonates exhibit elevated levels of IL-27 in the spleen and blood at resting state relative to adults (11, 12). To determine if IL-27 levels continue to rise during infection and how the cytokine may impact the host response, we established a murine model of neonatal sepsis. Neonatal pups were infected with *E. coli* O1:K1:H7 on day 4 of life. IL-27 gene expression was measured in the lungs, liver, spleen, kidneys, and brains of mice at 10 and 24 h following infection. These time points were chosen to span a critical window in the experimental model. These time points were chosen to span a critical window during infection. IL-27 gene expression varied with different tissues (Fig. 1A-B). While some infected pups expressed increased levels of IL-27p28 and EBI3 transcripts in all tissues examined, in several tissues, there were some pups that did not increase IL-27 gene expression (Fig. 1A-B). At the earlier 10 h time point, the most consistent increases in IL-27p28 and EBI3 expression were observed in the lungs, kidneys, and spleen of infected neonates (Fig. 1A). Surprisingly, the latter was the site of the greatest magnitude of increase in IL-27 expression (Fig. 1). At 24 h post-infection, more pups increased expression of IL-27 genes in all tissues except the liver; however, there were still some animals that maintained lower expression levels (Fig. 1B). To further analyze IL-27 systemically during infection, serum concentrations were measured by electrochemiluminescent immunoassay on days 1 and 2 post-infection. As shown in Figure 1C, IL-27 increased in circulation and peaked following the first day of infection in neonatal mice. While the mean IL-27 levels increased significantly more than three-fold, the population separated into higher and lower expressers similar to the gene expression data (Fig. 1C). A three-fold increase in IL-27 levels was maintained in infected pups relative to controls at day 2 post-infection, but at reduced overall magnitude (Fig. 1C). These data demonstrate that although IL-27 levels are higher at baseline in neonates relative to older populations, the levels continue to rise further during infection. Furthermore, the infected population of neonates at 24 h includes those expressing IL-27 genes and producing cytokine at higher levels, as well as those that better control IL-27 production. This could have implications on the progression and outcome of the infection.

**Figure 1:**
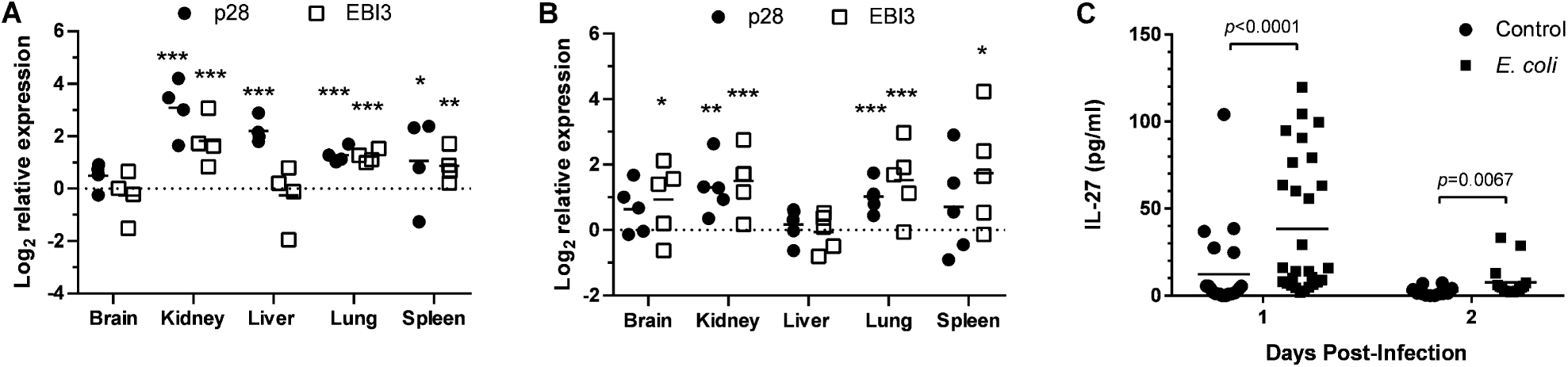
IL-27 levels rise during neonatal sepsis. Neonatal mice were subcutaneously inoculated with a target inoculum of ∼2×10^6^ CFUs/mouse of *E. coli* O1:K1:H7 or PBS as a control on day 3 or 4 of life. (**A, B**) At 10 (**A**) or 24 h (**B**) post-infection, the spleen, lung, kidney, and liver, and brain were harvested and RNA isolated. The expression of IL-27 p28 or EBI3 in infected tissues was determined relative to controls by real time PCR. Each symbol represents an individual animal finding with the mean for each group displayed. An individual experiment representative of two with similar results is shown.. (**A, B**) To assess IL-27 gene expression, non-parametric Mann Whitney U-Tests were performed on ΔCt values in control and infected samples for each tissue IL-27p28 and EIB3 at 10 and 24 h. The threshold for statistical significance was set to 0.05. Data are graphically represented as Log_2_ change in gene expression relative to control. Analyses revealed statistically significant changes in p28 gene expression in infected relative to control samples at 10 h in the kidney, liver, lung, and spleen (p<0.0001, p<0.0001, p<0.0001, and p=0.0449, respectively), and at 24 h in the kidney (p<0.0004) and lung (p<0.0001). Analyses revealed statistically significant differences in EBI3 gene expression in infected relative to control samples at 10 h in the kidney, lung, and spleen (p<0.0001, p<0.0001, and p=0.0083, respectively), and at 24 h in the brain (p=0.0321), kidney (p<0.0001), lung (p<0.0001), and spleen (p=0.0321). (**C**) Blood was collected from mice at day 1 or 2 post-infection and serum levels of IL-27 were measured by luminescent immunoassay. Statistical significance in the 95% confidence interval was determined using a Mann-Whitney U-Test; exact p values shown.

### Gr-1^+^ and F4/80^+^ cells are the most abundant IL-27 producers

We have previously shown that MDSCs and macrophages are the dominant cellular sources of IL-27 in neonatal mice in the absence of infection (11, 12). To determine cell types that contribute to the rising IL-27 levels during infection, we profiled cells in the blood and spleens by immunofluorescent labeling and flow cytometry. Both tissues are primary sites of infection and disseminated bacteria, as well as compartments with a significant population of myeloid cells. Our analysis evaluated IL-27 production in cells positive for Gr-1, F4/80, CD11c, and CD115. In the spleen and the blood, Gr-1^+^ cells were the most abundant cell type that produced IL-27 followed by a significant contribution from F4/80^+^ cells at 10 (Figs. 2A and B, 3A and B) and 24 h (Figs. 2A and D, 3A and D, S1, S2) post-infection. Surprisingly, there was no difference in the frequency of any IL-27-producing cell type in infected pups relative to controls in the spleen or the blood (Figs. 2, 3, S1, and S2). However, the mean fluorescent intensity (MFI) of IL-27 signal, indicative of the amount of IL-27 protein per cell, was increased in the spleen at 10 h in cells expressing all myeloid markers examined (Fig. 2C). CD115^+^ cells were associated with a substantial increase of nearly 100% during infection (Fig. 2C). Increased IL-27 expression was maintained at a significant level in CD11c^+^ cells at 24 h post-infection (Fig. 2E). Increased production in Gr-1^+^ and F4/80^+^ cells at 24 h was trending toward and nearly statistically significant (Fig. 2E). In the blood, only CD115^+^ and F4/80^+^ cells increased IL-27 production at either time point during infection, although these changes did not reach statistical significance (Fig. 3C and E). This analysis demonstrates that in the blood and spleen myeloid cells positive for Gr-1 and F4/80 are the most abundant producers of IL-27, while cells expressing all myeloid markers examined in the spleen likely contribute to the elevated IL-27 levels observed in infected neonates.

**Figure 2:**
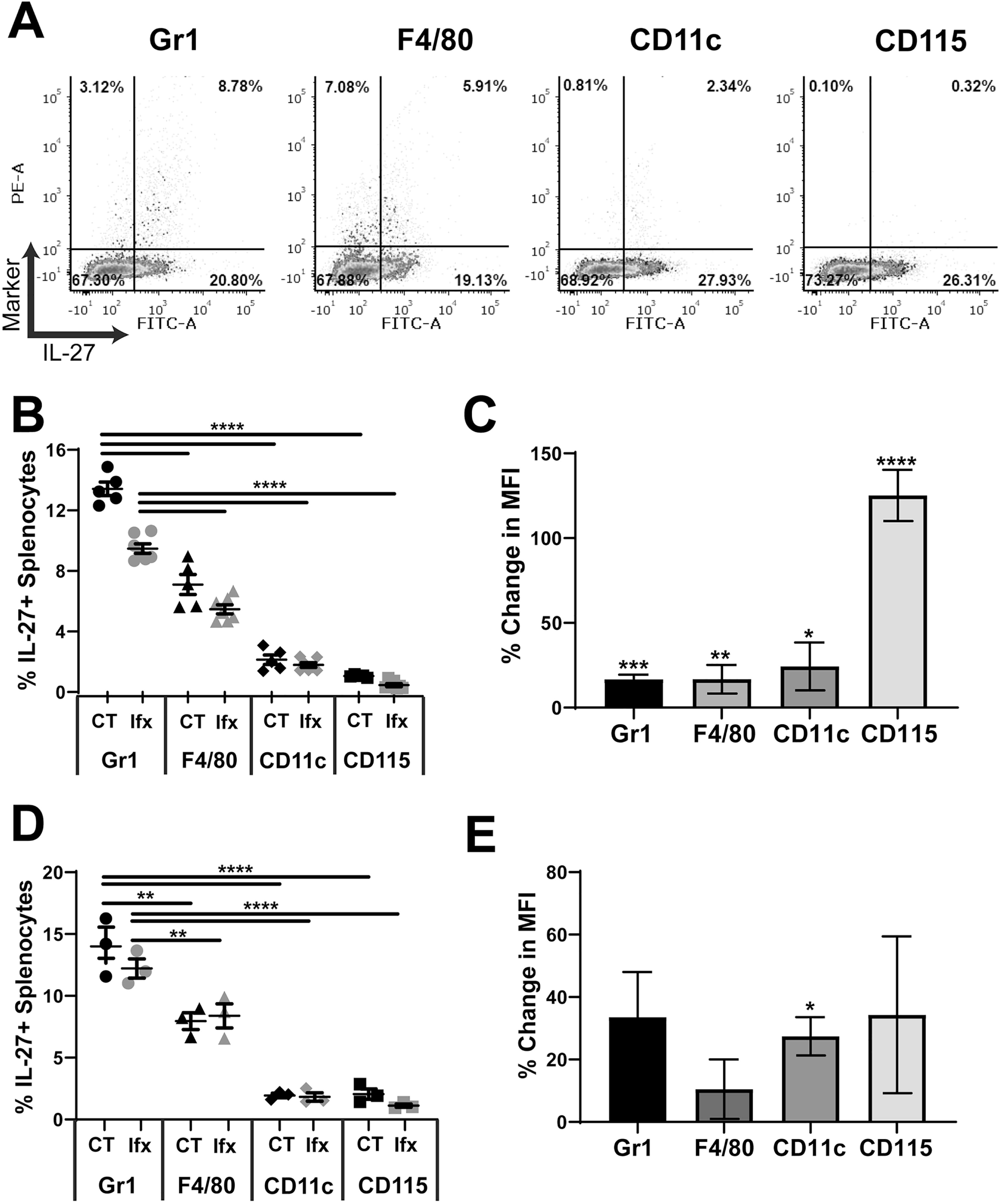
Cellular profiling of IL-27 producers in the spleen. Neonatal C57BL/6 (WT) mice were subcutaneously inoculated with a target inoculum of ∼2×10^6^ CFUs/mouse of *E. coli* O1:K1:H7 or PBS as a control on day 3 or 4 of life. At 10 or 24 h post-infection, mice were sacrificed and spleens were harvested. Single cell suspensions of splenocytes were immunolabeled for cell surface markers Gr-1, F4/80, CD11c, or CD115 and intracellular IL-27. Cells were analyzed by flow cytometry. Figures represent combined results from 2-3 independent experiments. **(A)** Representative dot plots of splenocytes labeled at 10 h post-infection for the indicated marker are shown. Cell surface markers are on the y-axis and IL-27 signal is on the x-axis represent. **(B)** The percentage double positive (upper right quadrant) of the population for each cell surface marker in control (CT) and *E. coli*-infected (Ifx) spleens at 10 **(B)** or 24 h **(D)** post-infection. **(C)** Percent change in mean fluorescence intensity (MFI) of FITC signal that corresponds to IL-27 protein in the double positive population for infected relative to control cells at 10 h **(C)** or 24 h **(E)** post-infection. **(B, D)** Statistical assessment was performed using a one-way ANOVA with Dunnett’s multiple comparison test. Means ± standard error are displayed. (**C, E**) Mean changes ± standard error in absolute values of MFI cell surface marker percentages at 10 h **(C)** and 24 h **(E)** post-infection in splenocytes were analyzed relative to a normalized baseline within the control groups using individual, unpaired t-tests for each cell surface marker. Asterisks indicate significant differences between infected and control splenocytes at 10 h **(C)** post-infection; Gr1 (*p*=0.0003), F4/80 (*p*=0.0027), CD11c (*p*=0.0237), and CD115 (*p*<0.0001). At 24 h **(E)** post-infection the asterisk indicates significance for CD11c (*p*=0.0111). Results shown for Gr-1 (*p*=0.08) and F4/80 (*p*=0.0689) were trending toward significance.

**Figure 3:**
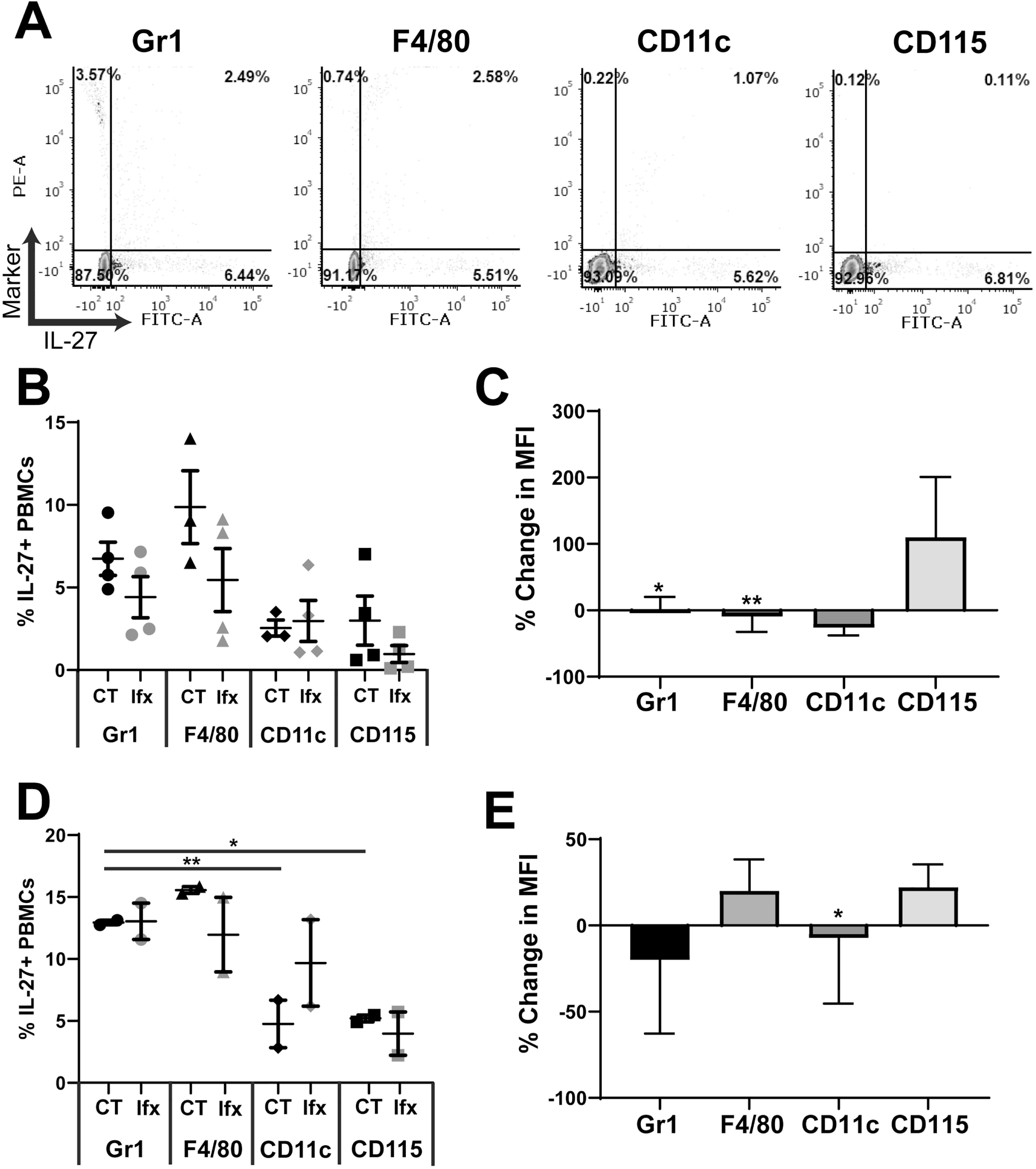
Cellular profiling of IL-27 producers in the blood. Neonatal C57BL/6 (WT) mice were subcutaneously inoculated with a target inoculum of ∼2×10^6^ CFUs/mouse of *E. coli* O1:K1:H7 or PBS as a control on day 3 or 4 of life. At 10 or 24 h post-infection, mice were sacrificed and blood was harvested and pooled for control and infected pups. PBMCs obtained by Ficoll density gradient centrifugation were immunolabeled for cell surface markers Gr-1, F4/80, CD11c, or CD115 and intracellular IL-27. Cells were analyzed by flow cytometry. Figures represent combined results from 2-3 independent experiments. **(A)** Representative dot plots of infected splenocytes at 10 h post-infection labeled for the indicated marker are shown. Cell surface markers are on the y-axis and IL-27 signal is on the x-axis represent. **(B)** The percentage double positive (upper right quadrant) of the population for each cell surface marker in control (CT) and *E. coli*-infected (Ifx) spleens at 10 h (**B**) or 24 h (**D**) post-infection. **(C)** Percent change in mean fluorescence intensity (MFI) of FITC signal that corresponds to IL-27 protein in the double positive population for infected relative to control cells at 10 h (**C**) or 24 h (**E**) post-infection. **(B, D)** Statistical assessment was performed using a one-way ANOVA with Dunnett’s multiple comparison test. Mean changes ± standard error are displayed. (**C, E**) Mean changes ± standard error in absolute values of MFI cell surface marker percentages at 10 h **(C)** and 24 h **(E)** post-infection in splenocytes were analyzed relative to a normalized baseline within the control groups using individual, unpaired t-tests for each cell surface marker. Asterisks indicate significant differences between infected and control splenocytes at 10 h **(C)** post-infection; Gr1 (*p*=0.0119) and F4/80 (*p*=0.0056). Results shown for CD11c (*p*=0.0850) at 10 h post-infection were trending toward significance. At 24 h **(E)** post-infection the asterisk indicates significance for CD11c (*p*=0.0345).

### IL-27 promotes mortality and poor weight gain during neonatal sepsis

We further investigated the impact of elevated IL-27 levels on survival during neonatal sepsis. Mice deficient for WSX-1 (KO) in the C57BL/6 background do not express a functional IL-27 receptor and cannot respond to the cytokine. Morbidity and mortality were monitored over 4 days of parallel infection in KO and wild-type (WT) mice. A striking improvement in survival was observed in the absence of IL-27 signaling (Fig. 4A). Infected pups gained weight at a level comparable to uninfected controls in the KO group (Fig. 4B). In contrast, infected WT pups lagged significantly behind control pups in weight gain, an indication of morbidity (Fig. 4B). When the change in weight was expressed relative to the control pups in each group, a highly significant improvement in weight gain was evident in WSX-1^-/-^ neonates (Fig. 4C). This has important implications in human neonatal sepsis. The highest level of serum IL-27 at 24 h post infection also correlates with a critical time period in disease progression (Fig. 1C and 4A). Pups that remain viable through 2 days most frequently remain viable through a 4 day infection (Fig. 4A). As such, the day 2 post-infection population is enriched for mice that are likely to remain viable through the duration of infection and may represent some survivor bias. Although we cannot definitively show that pups deceased at day 1 or pups viable at day 1 and deceased at day 2 had higher circulating levels of IL-27, the trends in IL-27 gene expression, serum levels, and mortality suggest the possibility that IL-27 levels are maintained at lower concentration in neonatal animals most likely to survive the infection (Fig. 1 and 4A). Collectively, these results indicate that IL-27 interferes with a protective host response.

**Figure 4:**
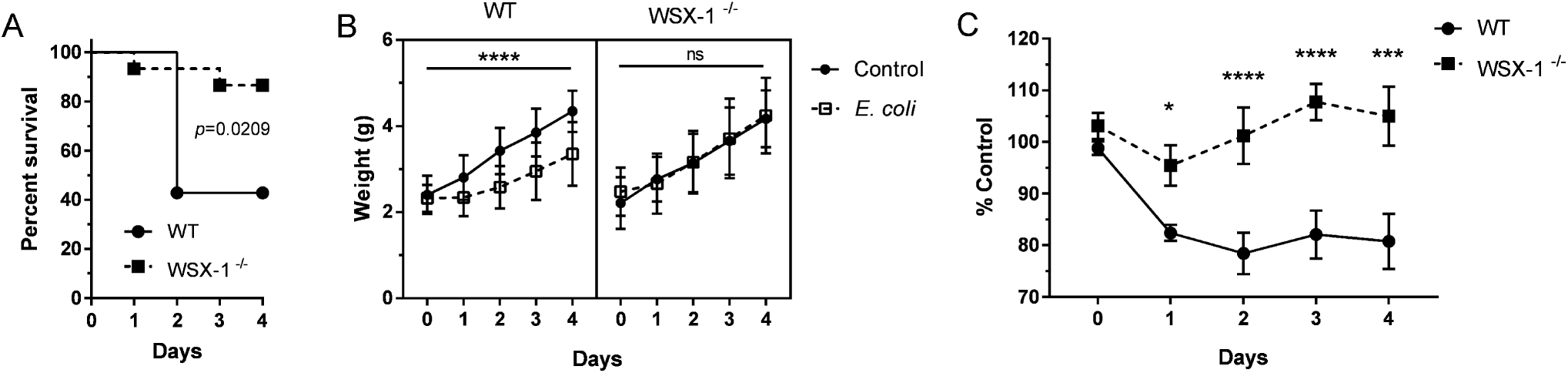
IL-27 promotes mortality and poor weight gain during neonatal sepsis. Neonatal C57BL/6 (WT) and WSX-1^-/-^ (KO) mice were subcutaneously inoculated with a LD_50_ dose of *E. coli* O1:K1:H7 or PBS as a control on day 3 or 4 of life and monitored daily for morbidity and mortality during infection. Three combined experiments for WT (n = 14) and KO (n=15) mice infected in parallel are shown. **(A)** Kaplan-Meier survival curves for WT and WSX-1^-/-^ mice over four days of infection. Statistical analysis was performed using the Mantel-Cox log rank test; exact p-value shown. **(B)** Recorded mean daily weight (g) ± SE for control and infected mice from WT (left) and WSX-1^-/-^ (right) in panel A above. A two-way ANOVA was used to determine statistical significance between control and *E. coli*-infected pups within the WT and WSX-1^-/-^ groups; an asterisk indicates p≤0.0001. (**C**) To compare WT and KO weight gain directly, the percent change for the infected pups relative to the control pups was represented for each day. A two-way ANOVA and Bonferroni multiple comparisons test was used to determine statistical significance between WT and KO groups; * indicates *p*≤0.0141, *** indicates *p*≤0.0001, **** *p*≤0.0003.

### IL-27 signaling opposes host clearance of bacteria during neonatal sepsis

To determine if improved survival in WSX^-/-^ mice is consistent with improved control of bacteria, we evaluated burdens in WT and KO mice 24 h post-infection. In the absence of IL-27 signaling, neonatal pups exhibited improved control of bacteria in the blood and all peripheral tissues examined (Fig. 5). To further explore mechanisms responsible for improved control of bacteria in WSX-1^-/-^ pups, we evaluated the ability of individual phagocytes to clear *E. coli in vitro*. The myeloid-restricted marker Ly6B.2 is highly expressed on neutrophils, inflammatory monocytes, and some populations of macrophages (40). Ly6B.2^+^ cells and bone marrow-derived macrophages (BMDMs) from WSX-1^-/-^ mice eliminated *E. coli* with increased efficiency early during infection (Fig. 5B and C). Similar results were obtained with F4/80^+^ cells isolated from WT and WSX-1 KO spleens (data not shown). TNF-α levels were lower in Ly6B.2^+^ cells from WSX-1^-/-^ pups during *in vitro* infection and marginally higher in BMDMs at 6 h only (Fig. 5D and E). Similar results were observed for IL-6 (data not shown). Improved bacterial clearance in WSX-1^-/-^ phagocytes in the absence of consistently higher levels of TNF-α, suggests that killing of bacteria is independent of proinflammatory cytokine production and may be a direct result of IL-27 signaling. We have previously reported that IL-27 opposes lysosomal acidification and trafficking in human macrophages with consequences to control of intracellular and extracellular bacterial growth (26, 27, 30, 31).

**Figure 5:**
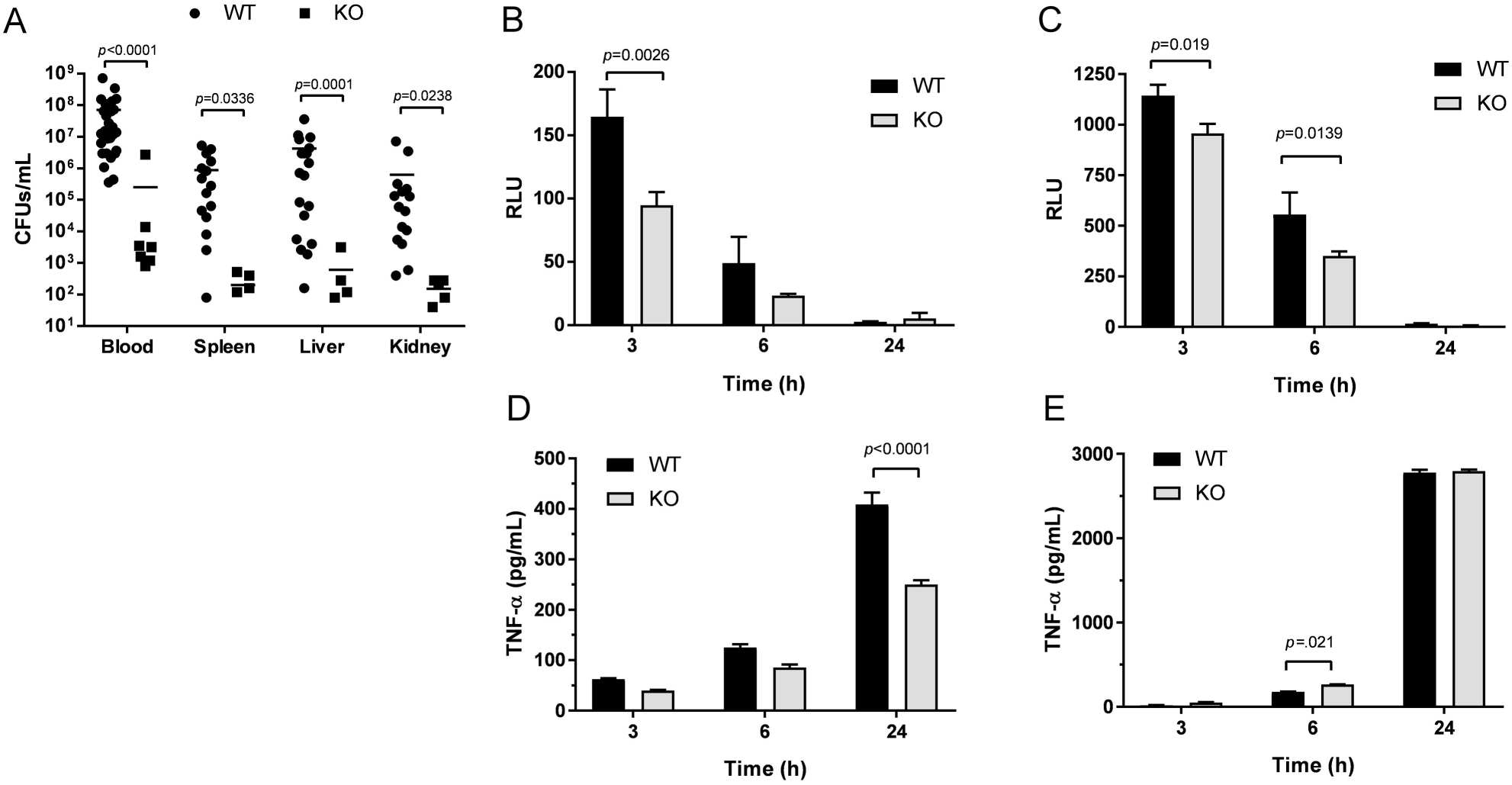
IL-27 signaling opposes host clearance of bacteria during neonatal sepsis. (**A**) Neonatal C57BL/6 (WT) and WSX-1^-/-^ (KO) mice were subcutaneously inoculated with a target inoculum of ∼2×10^6^ CFUs/mouse of *E. coli* O1:K1:H7 or saline as a control on day 3 or 4 of life. Peripheral tissues (spleen, liver, kidney) and blood were collected at 24 h post-infection and bacteria enumerated by standard plate counts. Colony forming units (CFUs)/mL for individual animals and experimental group means are shown for two combined experiments. Statistical significance in the 95% confidence interval was determined using a Mann-Whitney test; exact p-values shown. (**B, D**) Ly6B.2^+^ cells were isolated from the spleens of C57BL/6 (WT) and WSX-1^-/-^ (KO) neonatal mice. (**C, E**) Macrophages were derived from bone marrow progenitors obtained from C57BL/6 (WT) and WSX-1^-/-^ (KO) neonatal mice. (**B-E**) Cells were seeded in 96-well plates, and infected with luciferase-expressing *E. coli* O1:K1:H7 at an MOI of 50. After 1 h, the media was replaced with fresh that contained gentamicin (100 μg/mL). (**B, C**) Luminescence (RLU) was measured at 3 and 6 h post-infection. (**D, E**) Culture supernatants were collected at the indicated time and TNF-α measured by ELISA. (**B-E**) Mean findings ± SE for an individual experiment representative of multiple are shown. Statistical significance in the 95% confidence interval was determined using a two-way ANOVA and Bonferroni’s multiple comparisons test; exact p-values shown.

We next wanted to examine the kinetics of bacterial clearance in real time with a focus on the critical early phase of the infection. We infected WT and KO pups with luciferase-expressing *E. coli*, and longitudinally imaged individual mice over a 24-hour period. Consistent with CFU counts in harvested tissues, there was a robust difference in luminescent signal from KO pups. As a result, WT and KO mice could not be analyzed on the same luminescence scale. The drastic difference in luminescence resulted in oversaturation of signal in WT mice placed on the KO scale (Fig. S3A). Conversely, there was an absence of signal in KO mice placed on the WT scale at 10 and 24 h post-infection, further demonstrating the significant improvement in bacterial burdens in pups that cannot respond to IL-27 (Fig. 6A). Peak luminescent signal was observed at 10 h post-infection in KO pups, indicating that bacterial replication was controlled at this point in the infection (Fig. 6B and S3B). In contrast, the luminescence measured at 10 h in WT pups was increased relative to KO pups and continued to increase through 24 h (Fig. 6B and S3B). The final signal intensity at 24 h was nearly four orders of magnitude higher in WT pups (Fig. 6B,6C, and S3B). This real time imaging analysis also uncovered the brain as a site for high levels of bacteria in WT animals (Fig. 6C and S4). This finding was not unexpected since *E. coli* K1, including our strain, is a leading cause of neonatal meningitis (41, 42). However, the magnitude of difference between WT and KO pups was striking. In the absence of IL-27 signaling, there was a significant reduction in luminescent signal and CFUs in the brain (Fig. 6C and S4). Overall, the change in luminescence amongst mice correlated with actual CFUs in tissues and blood of WT and KO mice following imaging at 24 h (Fig. 6C).

**Figure 6:**
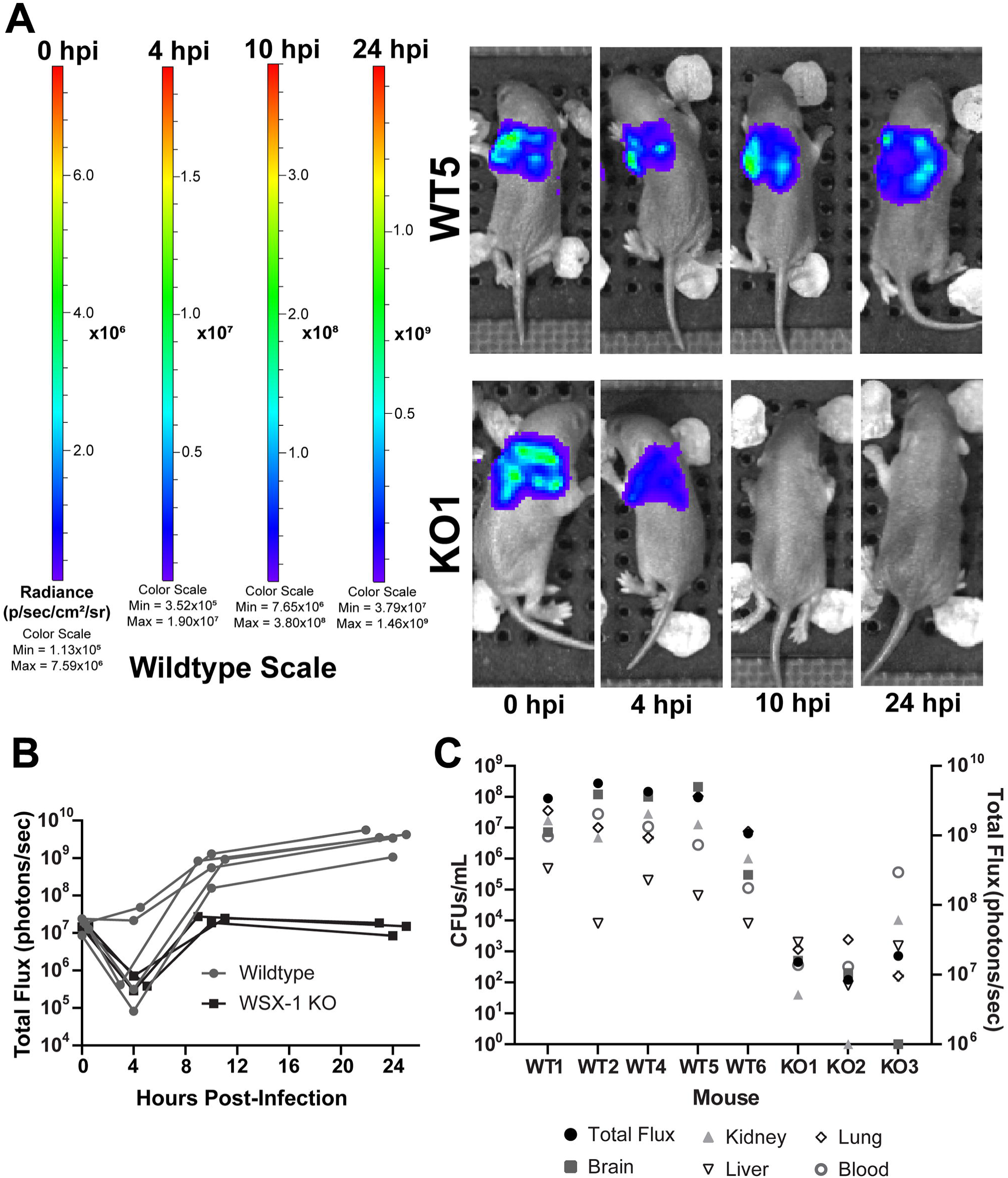
Intravital longitudinal imaging of the influence of IL-27 during neonatal sepsis. Neonatal C57BL/6 (WT) and WSX-1^-/-^ (KO) mice were subcutaneously inoculated with ∼2×10^6^ CFUs/mouse of luciferase-expressing *E. coli* O1:K1:H7 or PBS as a control on day 4 of life in parallel. The neonatal pups were imaged longitudinally on an IVIS SpectrumCT at 0, 4, 10, and 24 hours post-infection (hpi). Each mouse was tail-tattooed for individual identification during imaging. Data is the result of an independent experiment (WT n=5, KO n=3) representative of two with similar results. **(A)** Luminescent images of representative WT and KO mice at 0, 4, 10, and 24 hpi. The signal shown is on the WT scale. Colorimetric scale: low (minimum) signal is blue, intermediate signal is green, high (maximum) signal is red. **(B)** Total luminescent flux in photons/second for individual mice at 0, 4, 10, and 24 hpi. **(C)** At 24 hpi, mice were sacrificed and blood and peripheral tissues were collected for enumeration of bacteria by standard plate counts. Total luminescent flux (photons/second) and CFUs/mL from each tissue for individually infected mice are shown.

### WSX-1^-/-^ mice exhibit reduced levels of inflammation during infection

Failure to control bacterial replication promotes excessive and pathological inflammation during sepsis. As such, we evaluated gene expression levels of inflammatory cytokines in the spleens following one day of parallel infection in WT and KO neonates. This time point was chosen to evaluate pups during the critical phase; later time points would fail to include pups that succumb to infection and enrich the data set with findings from animals that exhibit improved outcomes. Gene expression levels were expressed relative to uninfected controls for WT and KO mice separately. WT levels of TNF-α, IL-1, and IL-6 increased robustly in WT pups following infection, while TNF-α and IL-6 expression were significantly reduced in WSX-1^-/-^ pups (Fig. 7A). Although there was a trend of reduced IL-1 expression in KO pups, this finding did not reach statistical significance (Fig. 7A). The levels of serum cytokines followed a similar pattern (Fig. 7B). IL-6 levels increased dramatically in infected WT pups and were maintained at a level two orders of magnitude lower in KO pups, while TNF-α and IL-1 levels were reduced approximately ten-fold (Fig. 7B). IL-6 levels are increased in patients with infectious complications and used clinically to provide a quantitative assessment of sepsis severity (43-45). Additionally, IL-6 levels correlate with the mortality rate in septic patients (46). The striking difference in IL-6 serum concentrations are reflective of peak illness in WT mice and a condition that is improved in mice that do not respond to IL-27 (47, 48).

**Figure 7:**
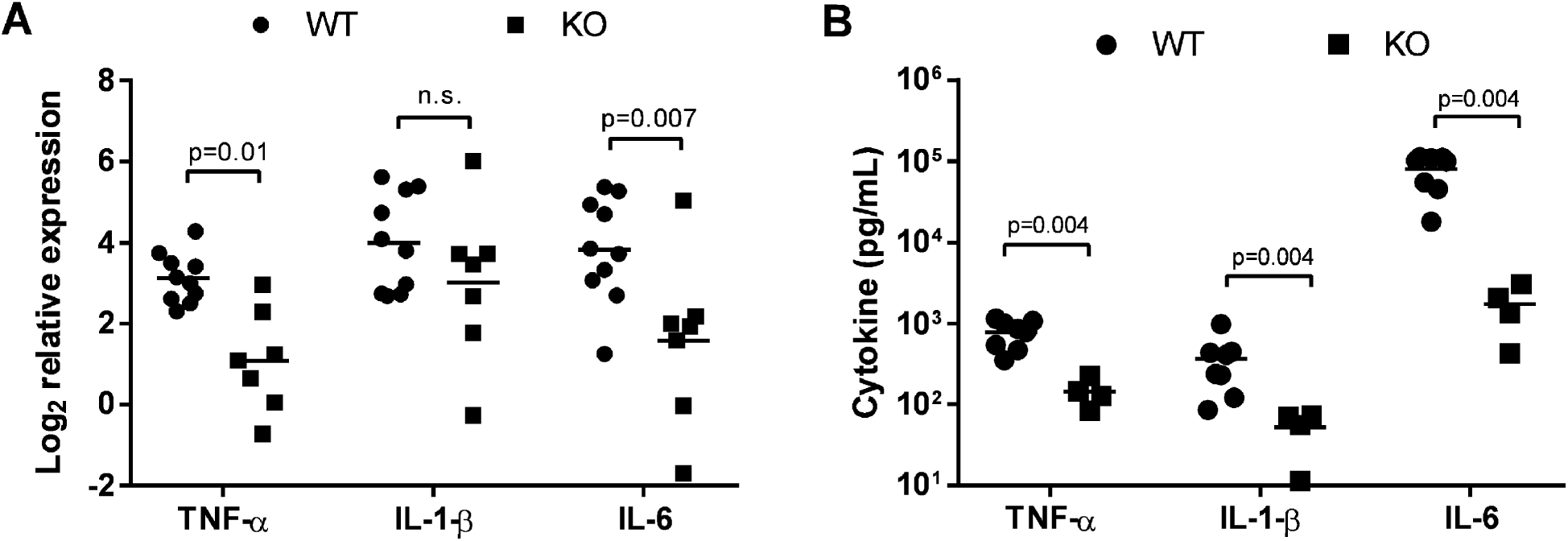
WSX-1^-/-^ mice exhibit reduced levels of inflammation during infection. Neonatal C57BL/6 (WT) and WSX-1^-/-^ (KO) mice were subcutaneously inoculated with a target inoculum of ∼2×10^6^ CFUs/mouse of *E. coli* O1:K1:H7 or PBS as a control on day 3 or 4 of life. (**A**) Spleens were harvested 1 day post-infection and RNA isolated. The expression of TNF-α, IL-1, and IL-6 were determined relative to controls by real time PCR. Individual animal findings and group means are shown for two combined experiments. Statistical significance in the 95% confidence interval was determined using individual t tests; exact p-values shown. **(B)** Blood was collected from mice at day 1 post-infection and serum levels of the indicated cytokine were measured by multiplex immunoassay. Individual animal findings and group means are shown for two combined experiments. Statistical significance in the 95% confidence interval was determined using a Mann-Whitney test; exact p-values shown.

## DISCUSSION

We have established elevated levels of the immune suppressive cytokine IL-27 at resting state in neonates, and we further demonstrate here that those levels continue to rise in most neonatal pups following infection that leads to sepsis. Our data regarding IL-27 serum levels are consistent with related findings in septic adult humans. Adult septic patients exhibit increased levels of IL-27 transcripts in whole blood and higher levels of serum cytokine (49). However, the absence of IL-27 data prior to infection limits our understanding of whether septic individuals are predisposed to higher IL-27 levels that constitute a risk factor for infection, or are elevated as a consequence of infection. Our model addresses this question specifically in neonates with littermates from inbred mice. Our findings have both parallels and contrasts with adult mice that become septic following caecal ligation and puncture (CLP) for which there is IL-27-related data. In these models, splenic IL-27 transcripts rise early peaking at 6-12 h, and protein levels are high in serum at 24 h (49-51). The greatest abundance of IL-27p28 transcripts at 6 h was found in the spleen and lungs (49-51). Our analysis identified the greatest increase in IL-27p28 and EBI3 transcripts during early infection in the lungs and kidneys of septic neonates. Later in the infection at 24 h, IL-27 transcripts were more widely increased across different tissues. In contrast to our neonatal data, adult mice maintain high levels of serum IL-27 through 72 h (49). Peak IL-27 serum levels at 24 h during neonatal infection may be influenced by the nature of the model and critical window for survival. We are the first to profile IL-27-producing cells in the blood and tissues during septic infection of any age group. Bosmann and colleagues depleted macrophages by clodronate treatment and observed a significant decline in IL-27 in the blood during endotoxic shock (51). Our analysis of IL-27-producing myeloid cells in the spleen and blood revealed Gr-1^+^ and F4/80^+^ cells as the most abundant source of cytokine. Neither of these surface markers is exclusive to a particular cell type. Gr-1 is expressed on MDSCs, some monocyte and macrophage populations, and at a high level on granulocytes (52-54). We recently described MDSCs as a significant source of IL-27 (12), and it is tempting to speculate that these cells are a significant source of rising levels during infection. F4/80 is expressed on monocytes, macrophages, and eosinophils (55, 56). An unexpected finding was that the frequency of these cells was not increased in septic pups. However, the mean fluorescent intensity of IL-27^+^ cells demonstrated that some cells elevated their level of cytokine production at different times during infection. This was true of all myeloid populations examined in the spleen. The increased IL-27 level in CD115^+^ cells was especially dramatic at 10 h post-infection. Our cellular profiling focused on myeloid cells, which are considered the dominant cellular sources of IL-27 (57). However, we cannot rule out other cellular sources that make a contribution to the overall levels of IL-27 produced during infection. Endothelial and epithelial cells have been reported to express both IL-27 subunits (15, 57). Contributions from these cells may help to explain the high levels of IL-27 expression observed in our model in the kidney, a tissue with extensive vasculature.

The presence of IL-27 in infected neonates in our system correlates with a significant increase in bacterial burdens and mortality. We have previously reported that IL-27 interferes with lysosomal acidification and trafficking in macrophages. This promotes increased growth of intracellular and extracellular pathogens (11, 27, 30, 31). Similarly, we recently reported that MDSC-derived IL-27 opposes control of *E. coli* by macrophages (12). In this report, we demonstrated improved clearance of bacteria by Ly6B.2^+^ myeloid cells and bone marrow derived macrophages from WSX-1^-/-^ pups. It is likely that the previously reported influence of IL-27 on lysosomal activity is directly responsible for enhanced killing of *E. coli* shown here (30, 31). Furthermore, since survival from sepsis in murine neonates does not depend on an intact adaptive immune system (58), the improved innate immune cell mediated clearance of bacteria in the absence of IL-27 is likely critical to the improved mortality in those neonatal pups. Lower levels of inflammatory cytokines in WSX-1-deficient neonates during infection is consistent with reduced bacterial burdens. Bacteria and bacterial-derived products drive the pathological inflammatory response during sepsis. Less inflammation in the absence of IL-27 may seem counterintuitive given many literature precedents, but our results suggest that the direct influence of IL-27 on bacterial killing by phagocytes is the dominant mechanism that dictates outcomes during neonatal sepsis. This implies that the greater numbers of circulating bacteria in WT pups drives an enhanced inflammatory response due to this negative influence of IL-27 on phagocyte clearance. Similarly, reduced concentrations of inflammatory cytokines and chemokines were found in the blood in adult mice given an IL-27 neutralizing antibody during endotoxic shock and in the lungs of WSX-1^-/-^ adult mice during a CLP-induced impairment of secondary bacterial challenge (49, 51). Fang and colleagues demonstrated that IL-27 neutralization reduced pulmonary inflammation in a mouse model of CLP-induced lung injury (59). It is also important to consider a possible effect of IL-27 on endothelial cells. IL-27 has been implicated in the endothelial dysfunction that occurs during cardiovascular pathology central to atherosclerosis by stimulating inflammatory cytokine and chemokine expression (60). Furthermore, IL-27 increased production of IL-6 and an inflammatory chemokine cascade in human endothelial cells (61, 62). This highlights the double-edge sword nature of IL-27. IL-27 has also been shown to activate an inflammatory response and suppress IL-10 production in monocytes (32). These cells would be expected to be significant players in the innate immune response during bacterial sepsis.

The intravital imaging analysis further supports the conclusion that IL-27 opposes bacterial clearance and allowed us to observe the rapid progression of dissemination that occurs in WT pups. To our knowledge, this is the first time bacterial sepsis has been imaged intravitally in neonatal mice. To this point, studies on sepsis in the context of LPS, group B streptococci, or *E. coli* in neonates have utilized confocal imaging of fixed tissue sections for analysis of bacterial load and inflammation (63-65). The presence of bacteria in the brain further validates our model as one that recapitulates findings clinically relevant in human neonates infected with *E. coli* K1 (41, 42). Our study represents a novel approach to understanding bacterial dissemination in a neonatal model relative to the host response, and drives home a direct association between IL-27 and severity of infection.

Improved infection control in adult mice lacking EBI3 or WSX-1 occurs during both *M. tuberculosis* and *P. aeruginosa* infections or CLP-induced peritonitis (21, 22, 49, 50). However, this improved infection outcome is in contrast to other studies that demonstrate elimination of IL-27 results in marked susceptibility to infection from *Trypanosoma cruzi, Trichuris muris, Leishmania major*, and *Toxoplasma gondii* (18, 20, 23, 66, 67). The differences in infection outcome relative to IL-27 suggests a microbe and Th1 vs Th2-dependent mechanism of immunity, as well as a potential threshold of IL-27 production necessary to modulate proper immunity. Although IL-27 may serve a beneficial role in the balance of inflammatory response, in different infectious contexts, over or underproduction of this cytokine may result in immune dysregulation and pathogen expansion.

There were some limitations to our study. Overall, the number of cells that could be obtained from the blood and spleen were limited. As a result, we could not perform more extensive profiling of IL-27-producing cells. As mentioned previously, non-myeloid cells may contribute to the total IL-27 levels and may even be undervalued in that regard. Additionally, we have not developed an approach that allows for blood sampling from viable neonates. As such, we were unable to follow each pup for IL-27 levels and subsequent bacterial burdens or viability versus mortality. The technical ability to perform this level of analysis would further strengthen our conclusions.

In summary, our results suggest that elevated levels of IL-27 early in life predispose to impaired control of pathogen burden further compounded by continued increases in circulating levels of IL-27 during sepsis. These findings have enormous translational potential. On the diagnostic front, IL-27 levels in circulation may predict susceptibility to septic infection and related outcomes. Similarly, IL-27 levels may predict outcomes and guide initiation of antibiotic therapy in neonates that appear ill. Indeed, IL-27 has been proposed as a biomarker for neonatal sepsis (37). IL-27 antagonism may also offer therapeutic potential. Our results predict reducing IL-27 levels will promote bacterial clearance, improve host response, and reduce mortality. This approach may have prophylactic value for populations at increased risk in addition to a post-infection therapy. Currently, the only available treatment options to combat bacterial sepsis are antibiotics and supportive care (68). IL-27 may represent a targeted adjunctive therapy to augment the efficacy of antibiotics to improve survival and infection-related outcomes in neonates.

## FIGURE LEGENDS

**Supplemental Figure 1:Cellular profiling of IL-27 producers in the spleen.** Neonatal C57BL/6 (WT) mice were subcutaneously inoculated with a target inoculum of ∼2×10^6^ CFUs/mouse of *E. coli* O1:K1:H7 or PBS as a control on day 3 or 4 of life. At 10 or 24 h post-infection, mice were sacrificed and spleens were harvested. Single cell suspensions of splenocytes were immunolabeled for cell surface markers Gr-1, F4/80, CD11c, or CD115 and intracellular IL-27. Cells were analyzed by flow cytometry. Results from control pups at 10 h (**A**) or 24 h (**B**) are shown. (**C**) Results from infected pups at 24 h; 10 h dot plots were shown in Figure 2.

**Supplemental Figure 2:Cellular profiling of IL-27 producers in the blood.** Neonatal C57BL/6 (WT) mice were subcutaneously inoculated with a target inoculum of ∼2×10^6^ CFUs/mouse of *E. coli* O1:K1:H7 or PBS as a control on day 3 or 4 of life. At 10 or 24 h post-infection, mice were sacrificed and blood was collected. Single cell suspensions of PBMCs were immunolabeled for cell surface markers Gr1, F4/80, CD11c, or CD115 and intracellular IL-27. Cells were analyzed by flow cytometry. Results from control pups at 10 h (**A**) or 24 h (**B**) are shown. (**C**) Results from infected pups at 24 h; 10 h dot plots were shown in Figure 3.

**Supplemental Figure 3:Intravital longitudinal imaging of the influence of IL-27 during neonatal sepsis requires separate scales for WT and WSX-1^-/-^ mice.** Neonatal C57BL/6 (WT) and WSX-1^-/-^ (KO) mice were subcutaneously inoculated with a target inoculum of ∼2×10^6^ CFUs/mouse of luminescent *E. coli* O1:K1:H7 and imaged longitudinally on an IVIS SpectrumCT at 0, 4, 10, and 24 hours post-infection (hpi). Each mouse was tail-tattooed for individual identification during imaging. At 24 hpi, mice were sacrificed, blood was collected for serum and bacterial burdens, and peripheral tissues were homogenized for bacterial burdens. **(A and B)** Data is the result of one independent experiment. n = 5 and 3 for wildtype and WSX-1 deficient mice, respectively. **(A)** Longitudinal luminescence images of representative wildtype and WSX-1 deficient mice at 0, 4, 10, and 24 hpi. Signal is on the KO scale. Colorimetric scale: low (minimum) signal is blue, intermediate signal is green, high (maximum) signal is red. **(B)** Pooled bacterial luminescence signal (total flux) represented as photons/second for wildtype and WSX-1 deficient mice at 4, 10, and 24 hpi. Black circle symbols represent each individual mouse in wildtype infections, black square symbols represent each individual mouse in knockout infections. Statistical analysis of **(B)** was carried out using a nonparametric Mann-Whitney U test, median with interquartile range displayed. * = p≤0.05.

**Supplemental Figure 4:Intravital imaging reveals the brain as an organ associated with high bacterial burdens during sepsis.** Neonatal C57BL/6 (WT) and WSX-1^-/-^ (KO) mice were subcutaneously inoculated with a target inoculum of ∼2×10^6^ CFUs/mouse of luminescent *E. coli* O1:K1:H7 and imaged longitudinally on an IVIS SpectrumCT at 24 hours post-infection (hpi). Images are representative of 2 individual experiments. **(A)** Representative CT images of two WT mice with bacterial infection in their brains. Signal is on the wildtype (WT) scale. **(B)** Representative CT images of one KO mouse with bacterial infection in the brain. Signal is on the knockout (KO) scale. Perspective (x, y, z), coronal (x, y), and transaxial (x, z) views are shown from 3D CT images for both WT and KO mice. Colorimetric scale: low (minimum) signal is blue, intermediate signal is green, high (maximum) signal is red. White arrowheads directly point to burdens in brains.

