## Supplemental Figures 1-4 for "Elevated levels of interleukin-27 in early life compromise protective immunity during neonatal sepsis"

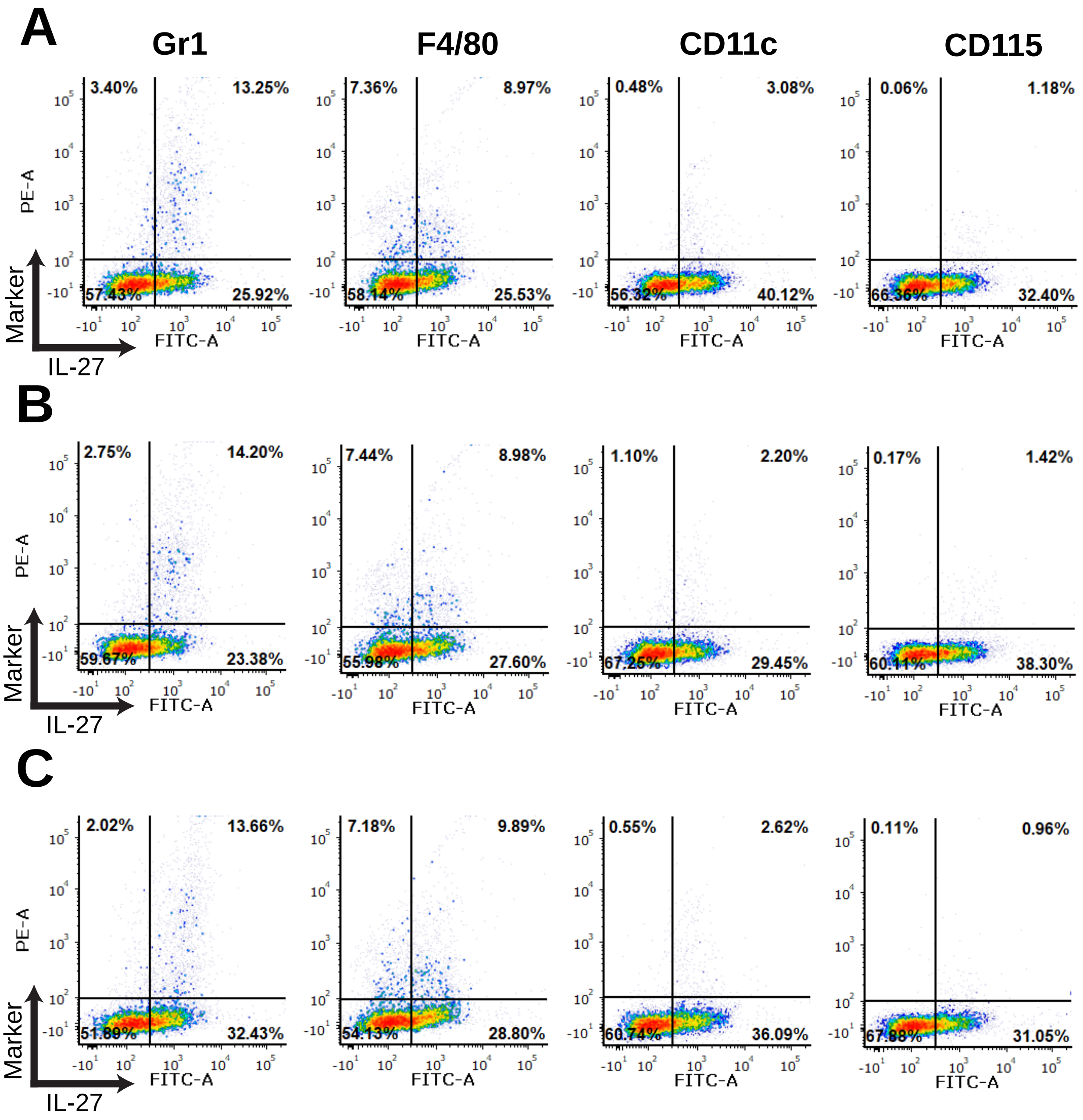
A
B
C
Gr1
F4/80
CD11c
CD115
PE-A
FITC-A
Marker
IL-27
3.40%
13.25%
57.43%
25.92%
7.36%
8.97%
58.14%
25.53%
0.48%
3.08%
56.32%
40.12%
0.06%
1.18%
66.36%
32.40%
2.75%
14.20%
59.67%
23.38%
7.44%
8.98%
55.98%
27.60%
1.10%
2.20%
67.25%
29.45%
0.17%
1.42%
60.11%
38.30%
2.02%
13.66%
51.89%
32.43%
7.18%
9.89%
54.13%
28.80%
0.55%
2.62%
60.74%
36.09%
0.11%
0.96%
67.88%
31.05%

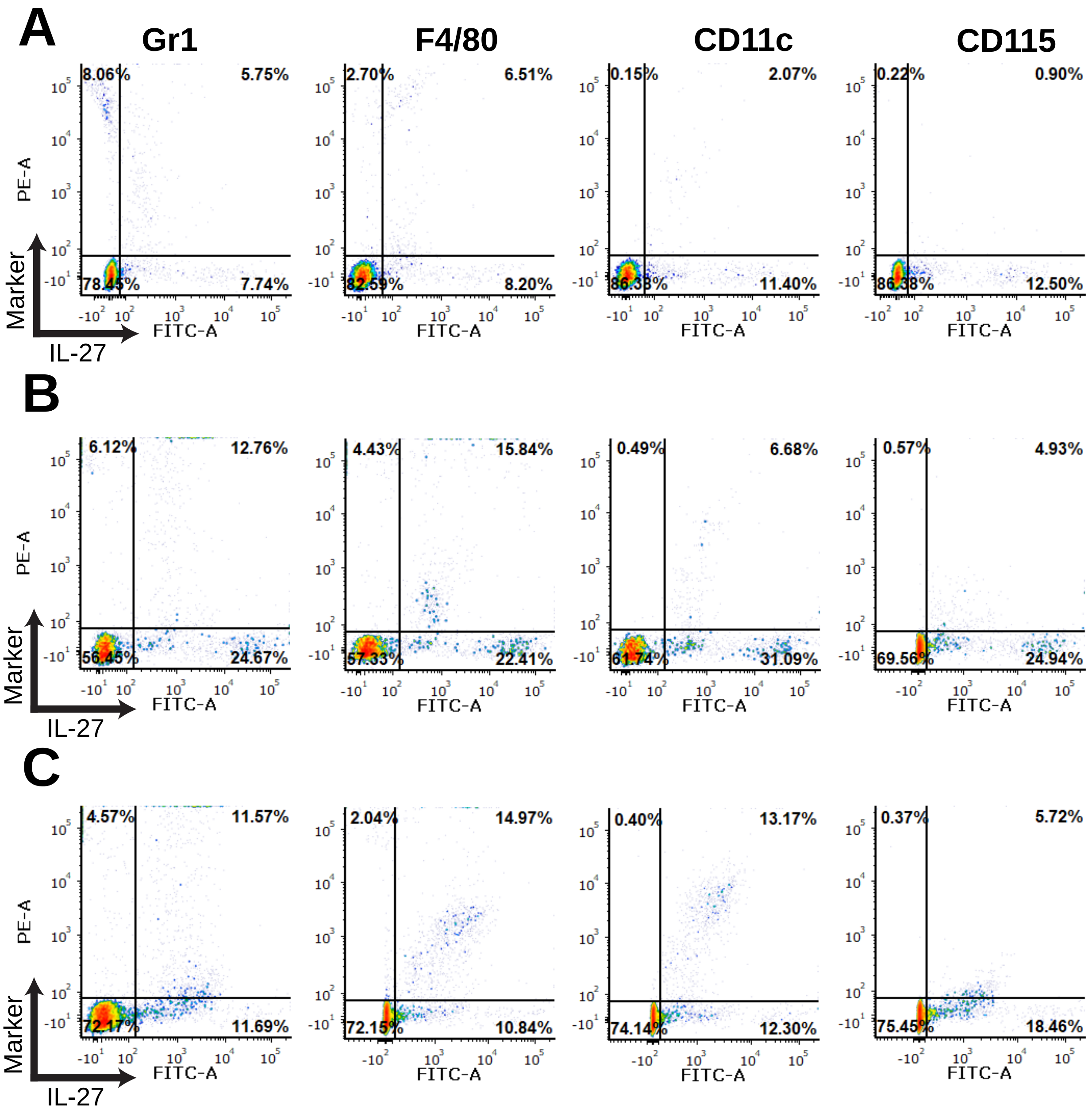
A
Gr1
F4/80
CD11c
CD115
8.06%
5.75%
78.45%
7.74%
2.70%
6.51%
82.59%
8.20%
0.15%
2.07%
86.38%
11.40%
0.22%
0.90%
86.38%
12.50%
B
6.12%
12.76%
56.45%
24.67%
4.43%
15.84%
57.33%
22.41%
0.49%
6.68%
61.74%
31.09%
0.57%
4.93%
69.56%
24.94%
C
4.57%
11.57%
72.17%
11.69%
2.04%
14.97%
72.15%
10.84%
0.40%
13.17%
74.14%
12.30%
0.37%
5.72%
75.45%
18.46%
PE-A
FITC-A
Marker
IL-27

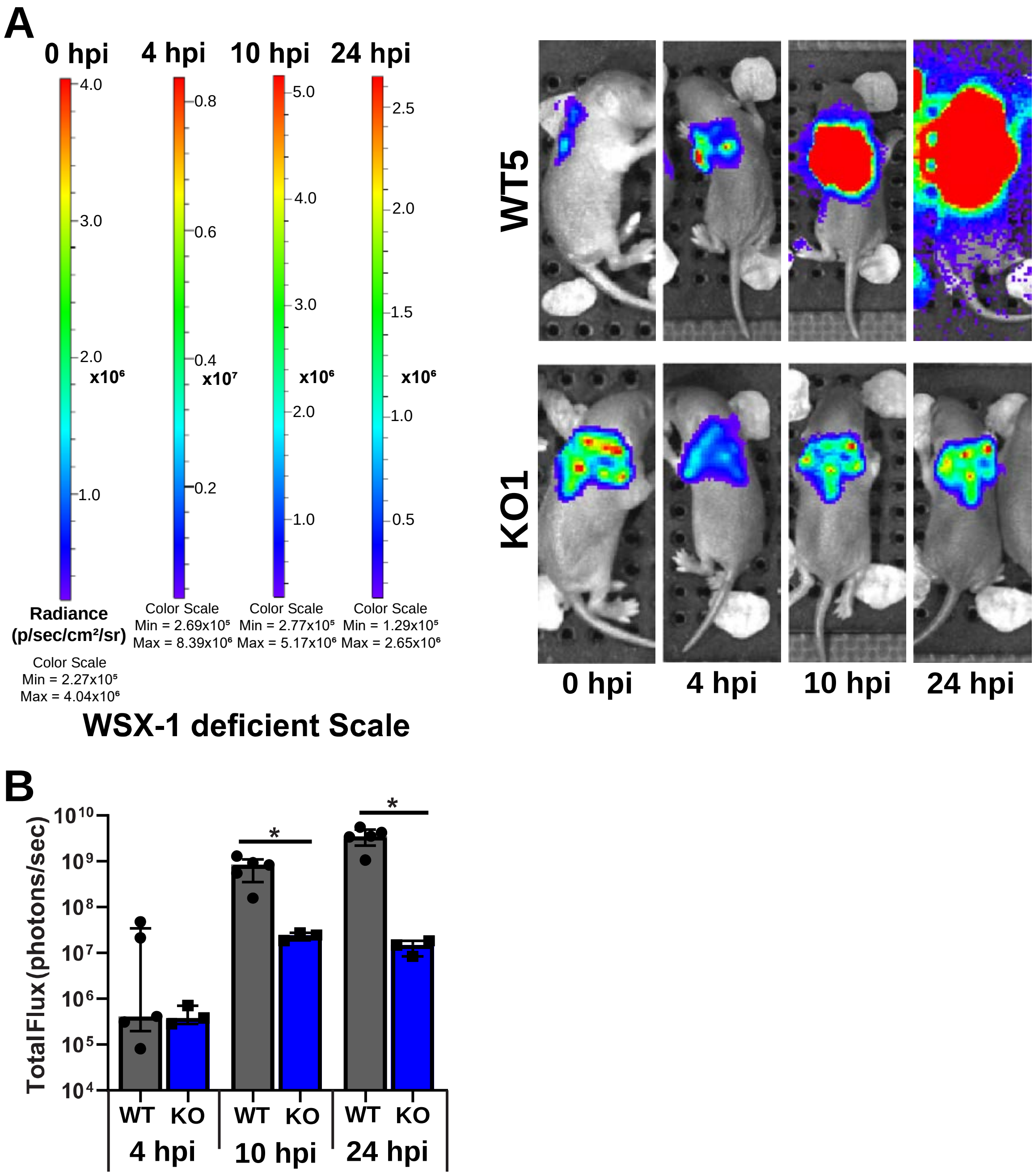

A
0 hpi
4 hpi
10 hpi
24 hpi
4.0
3.0
2.0
1.0
x10⁶
0.8
0.6
0.4
0.2
x10⁷
5.0
4.0
3.0
2.0
1.0
x10⁶
2.5
2.0
1.5
1.0
0.5
x10⁶
Radiance (p/sec/cm²/sr)
Color Scale Min = 2.27x10⁵ Max = 4.04x10⁶
Color Scale Min = 2.69x10⁵ Max = 8.39x10⁶
Color Scale Min = 2.77x10⁵ Max = 5.17x10⁶
Color Scale Min = 1.29x10⁵ Max = 2.65x10⁶
WSX-1 deficient Scale
WT5
KO1
0 hpi
4 hpi
10 hpi
24 hpi
B
Total Flux (photons/sec)
10¹⁰
10⁹
10⁸
10⁷
10⁶
10⁵
10⁴
*
*
WT
KO
WT
KO
WT
KO
4 hpi
10 hpi
24 hpi

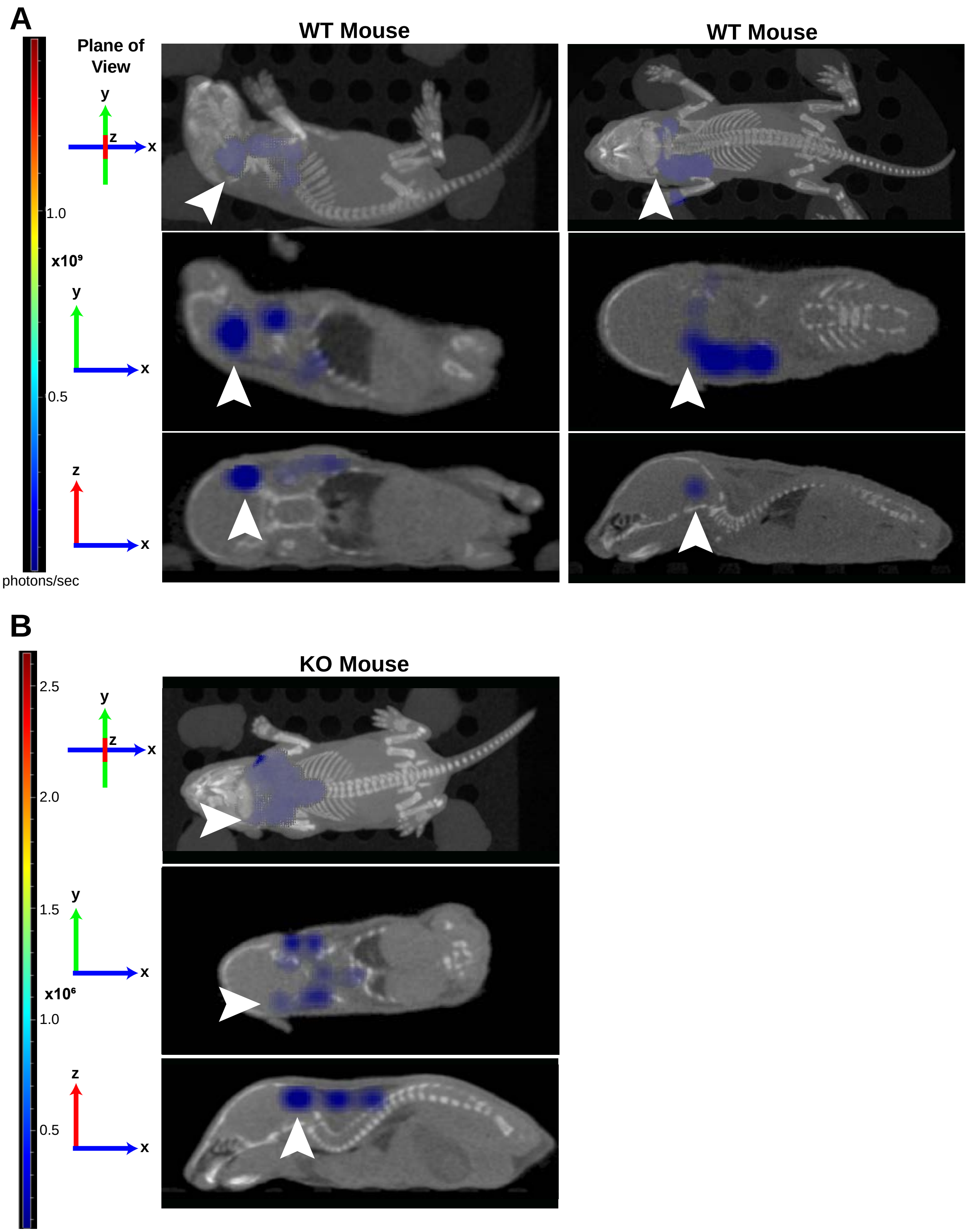

A
WT Mouse
WT Mouse
Plane of View
y
z
x
1.0
x10⁹
0.5
photons/sec
B
KO Mouse
2.5
2.0
1.5
x10⁶
1.0
0.5

**Supplemental Figure 1: Cellular profiling of IL-27 producers in the spleen.** Neonatal C57BL/6 (WT) mice were subcutaneously inoculated with a target inoculum of ~2x$10^6$ CFUs/mouse of *E. coli* O1:K1:H7 or PBS as a control on day 3 or 4 of life. At 10 or 24 h post-infection, mice were sacrificed and spleens were harvested. Single cell suspensions of splenocytes were immunolabeled for cell surface markers Gr-1, F4/80, CD11c, or CD115 and intracellular IL-27. Cells were analyzed by flow cytometry. Results from control pups at 10 h (**A**) or 24 h (**B**) are shown. (**C**) Results from infected pups at 24 h; 10 h dot plots were shown in Figure 2.
